# Visualization of ligand-induced transmembrane signalling in the full-length human insulin receptor

**DOI:** 10.1101/207928

**Authors:** Theresia Gutmann, Kelly H. Kim, Michal Grzybek, Thomas Walz, Ünal Coskun

**Affiliations:** Paul Langerhans Institute Dresden of the Helmholtz Zentrum München at the University Hospital and Faculty of Medicine Carl Gustav Carus of TU Dresden, TU Dresden, Fetscher Str. 74, 01307 Dresden, Germany; German Center for Diabetes Research (DZD e.V.), Ingolstädter Landstraße 1, 85764 Neuherberg, Germany; Laboratory of Molecular Electron Microscopy, The Rockefeller University, New York, NY 10065, USA

## Abstract

Using glycosylated full-length human insulin receptor reconstituted into lipid nanodiscs, we show that insulin binding to the dimeric receptor converts its ectodomains from an inverted U-shaped to a T-shaped conformation. This unprecedented structural rearrangement of the ectodomains propagates to the transmembrane domains, which are well separated in the inactive conformation, but come together upon insulin binding, allowing autophosphorylation of the cytoplasmic kinase domains.

## MAIN TEXT

The insulin receptor (IR) is a receptor tyrosine kinase involved in regulating glucose, protein, and lipid metabolism and growth. Its dysfunction is linked to severe pathologies, such as diabetes mellitus, cancer, and Alzheimer’s disease (1). IRs are unique in that they exist as preformed (αβ)_2_ homodimers at the cell surface (**Fig. 1a**), excluding ligand-induced dimerization, used by other receptor tyrosine kinases, as their activation mechanism. Our understanding of IRs derives from structures of receptor fragments (2–7), most notably an unliganded incomplete ectodomain (ECD) (7) and a truncated microreceptor–insulin complex (3).

**Figure 1.**
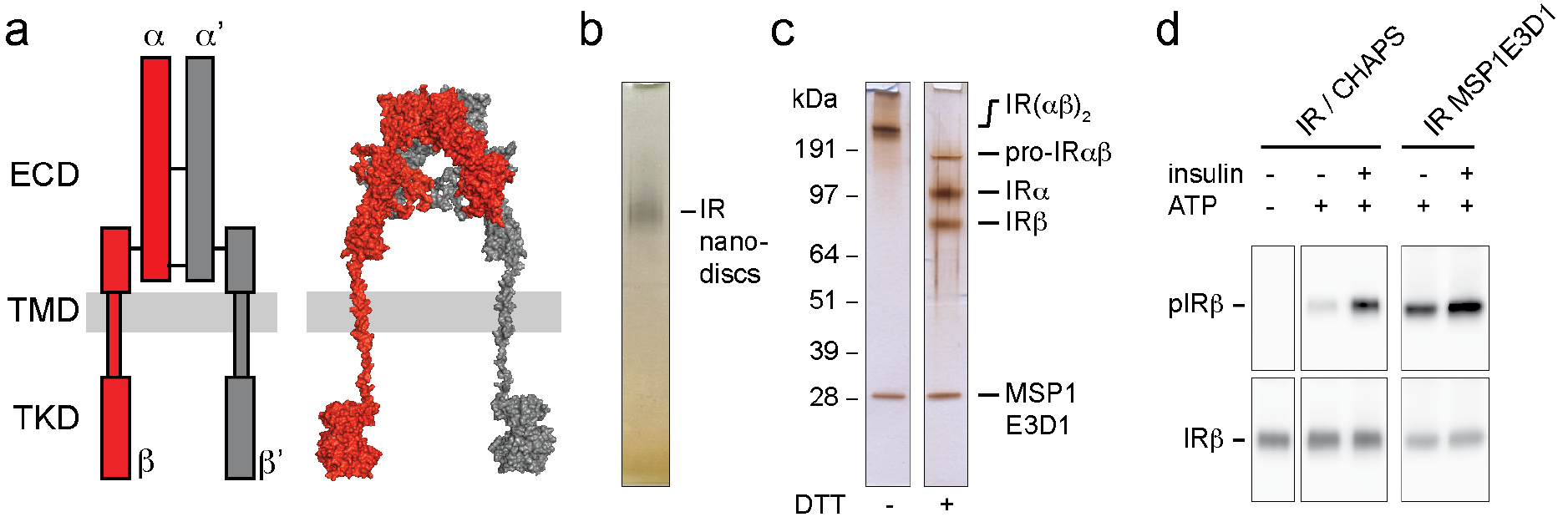
IR reconstitution into nanodiscs and activity assay. (**a**) Schematic cartoon showing the ectodomain (ECD), transmembrane domain (TMD) and tyrosine kinase domain (TKD) (left) and structural model (right) of the full-length human (αβ)_2_ IR. (**b**) Silver-stained native PAGE of glycosylated full-length human IR reconstituted into MSP1E3D1 nanodiscs. (**c**) SDS-PAGE of nanodisc-embedded IR under non-reducing (-DTT) and reducing conditions (+DTT). (**d**) Activity assay showing that both CHAPS-solubilized and nanodisc-reconstituted IRs are autophosphorylated upon insulin binding. IRα, IR α subunit; IRβ, IR β subunit; pIRβ, phosphorylated IRβ; IR(αβ)_2_, mature IR; pro-IRαβ, unprocessed intracellular form of IR. Experimental details are provided in **Experimental Procedures** and **SI Figures 12-14.**

The mechanism how insulin binding to the extracellular domains (ECDs) is transmitted by the transmembrane domains (TMDs) across the membrane to the cytoplasmic tyrosine kinase domains (TKDs) is unknown (8). To address this question, we produced recombinant full-length human receptor in HEK293 cells and purified it in CHAPS to near-homogeneity by a single affinity-chromatography step. The receptor was then reconstituted into nanodiscs with membrane scaffold proteins (MSPs) and a ternary lipid mixture (**Fig. 1b, c**). An insulin-stimulated kinase-activation assay demonstrated that the detergent-solubilised and nanodisc-embedded IRs were biologically active (**Fig. 1d**).

Full-length IR was first reconstituted with MSP1E3D1, which forms nanodiscs of ~12 nm diameter (9), and the resulting structures were imaged by negative-stain electron microscopy (EM). Although heterogeneous, many particles showed circular shapes representing nanodiscs, from which additional densities extended (**SI Fig. 1a**). About 10,000 interactively selected particles were subjected to *K*-means classification into 200 classes (**SI Fig. 1b**) to quantify particle populations (see below), and to iterative stable alignment and clustering (ISAC) (**SI Fig. 1c**) to produce averages with the crispest features (for classification details, see **SI Table 1**). Some ISAC averages revealed an L-shaped density extending from a nanodisc (**SI Fig. 1c**), presumably representing unprocessed monomeric IRs. Since monomeric IRs do not exist at the cell surface, they will not be discussed further. Many ISAC averages showed an inverted U-shaped density that either extended from a single nanodisc (20%; see Experimental Procedures for estimation of particle populations) (**Fig. 2a**, top panels) or connected two nanodiscs (80%) (**Fig. 2a**, bottom panels). Based on a previous negative-stain EM study (10) and available crystal structures (2,7), the U-shaped density represents the dimeric IR ECD. Furthermore, since most IR dimers were reconstituted into two separate nanodiscs, the two TMDs must generally be well separated from each other in unliganded receptors.

**Figure 2.**
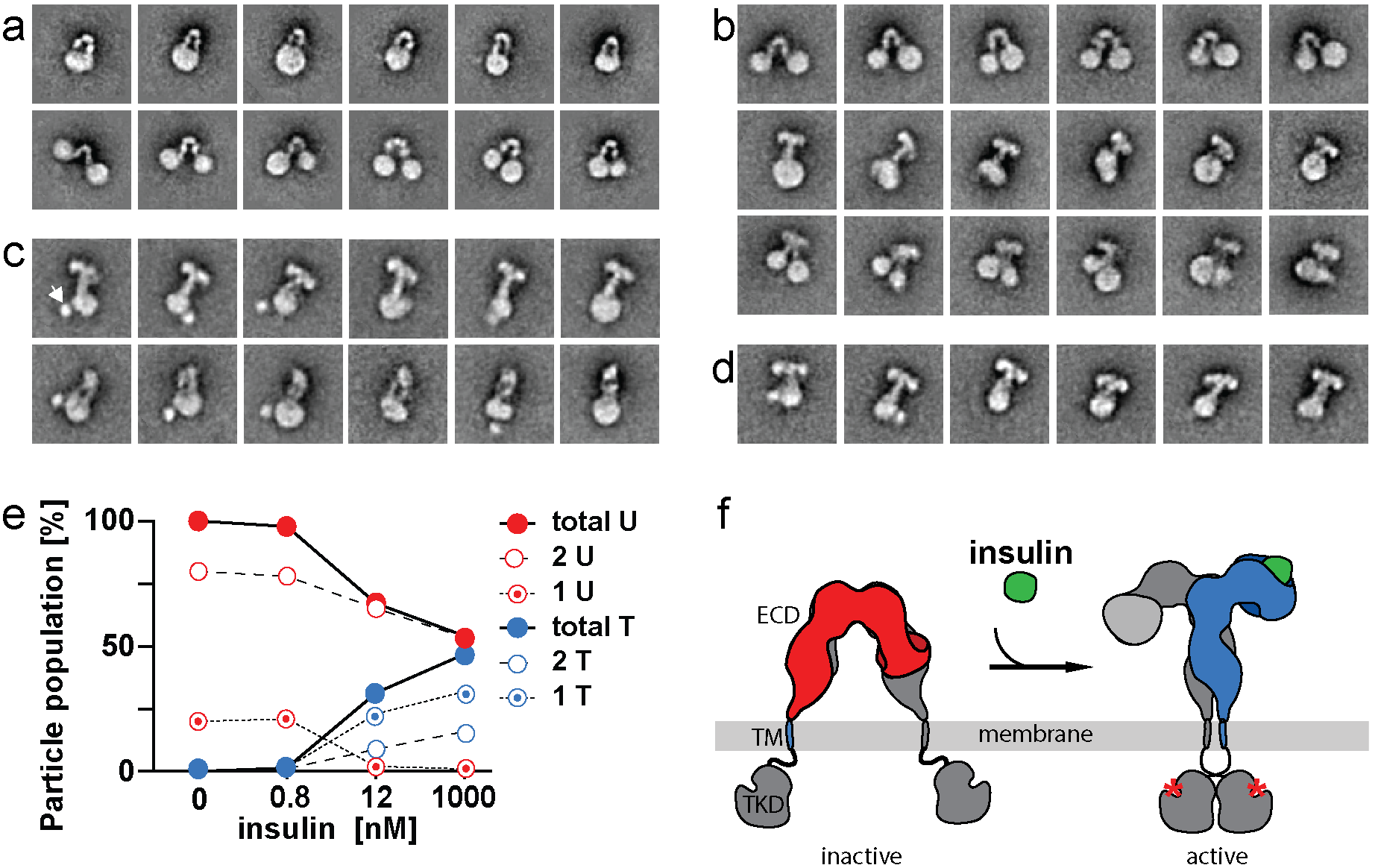
Selected EM averages of IRs reconstituted into MSP1E3D1 nanodiscs and ligand-induced IR activation. (**a**) In absence of insulin, IRs reconstituted into one (top panels) or two nanodiscs (bottom panels) and adopted a U-shaped conformation. (**b**) Upon insulin addition, all IRs in a single nanodisc adopted the T-shaped conformation (middle panels), whereas IRs in two nanodiscs adopted either the U-shaped (top panels) or T-shaped conformation (bottom panels). (**c**) In presence of insulin, all IRs reconstituted into a single nanodisc. Most of them adopted a T-shaped conformation (top panels) while some showed an II-shaped conformation (likely representing a different view of T-shaped IR) (bottom panels). The first three averages show a globular density that may represent dimerized TKDs (arrow). (**d**) Upon insulin addition, all IRs showed a T-shaped conformation. Side length of individual panels: 47.7 nm. (**e**) Quantification of IR populations with a particular ECD conformation at given insulin concentrations. (**f**) Cartoon illustrating ligand (green)-driven conformational change in the ECDs and coupling to the TMDs with concomitant activation by autophosphorylation (red asterisks) of the TKDs.

Images of IR reconstituted in presence of insulin showed that all receptors were incorporated into a single nanodisc (**SI Fig. 2**). All averages showed a dramatic conformational change from the U-shaped conformation predominantly into a T-shaped conformation (~72%) (**Fig. 2c**, top panels). A small fraction showed II-shaped densities (26%), presumably representing a different view of the T-shaped conformation (**Fig. 2c**, bottom panels). Furthermore, since all IR dimers are in a single nanodisc, the two TMDs must be in close proximity and presumably interact with each other in ligand-bound receptors.

To confirm that the observed conformational change also occurs in membrane-embedded IRs, we used IRs reconstituted into nanodiscs in absence of insulin (**Fig. 2a**) and exposed them to 1 insulin (**SI Fig. 3**). Averages still revealed IRs in the U-shaped conformation connecting two nanodiscs (**Fig. 2b**), but these constituted a smaller fraction (~53%). Strikingly, all IRs that were reconstituted into a single nanodisc appear to have undergone the conversion from U- to T-shaped conformation upon insulin binding. In addition, averages now also showed T-shaped IRs extending from two nanodiscs (~16%) (**Fig. 2b**). From the previous results, ligand binding to IRs reconstituted into two nanodiscs should convert their ECDs into a T-shaped conformation, which, in turn, would exert a driving force to bring the TMDs together. This force will tend to bring the two nanodiscs together, which is seen in some of the averages (**Fig. 2b**, bottom panels), and nanodiscs connected by a T-shaped IR in general appear to be closer to each other than ones connected by a U-shaped IR (**Fig. 2b**, compare top and bottom panels). The resulting juxtaposition of the two nanodiscs may only occasionally lead to fusion, allowing the TMDs to interact, while in most cases the two nanodiscs will likely remain separate, preventing the TMDs from interacting.

Exposure of IR reconstituted into nanodiscs in absence of insulin or to lower insulin concentrations, 0.8 nM (**SI Fig. 4**) and 12 nM (**SI Fig. 5**), showed that the insulin binding-induced conversion of the ECD from U- to T-shaped conformation is dose-dependent (**Fig. 2e**). Also, consistent with all results so far, analysis of IR reconstituted into nanodiscs in presence of insulin and exposed further to 1 μΜ insulin (**SI Fig. 6**) showed that all IR dimers incorporated into a single nanodisc and adopted a T-shaped conformation (**Fig. 2d**).

We repeated these experiments with MSP1D1, which forms smaller nanodiscs of ~10 nm diameter (9). As expected, in absence of insulin, all receptors were in the U-shape conformation, but because of the smaller nanodisc size, all IRs were in two nanodiscs, strengthening the notion that TMDs in unliganded receptors do not interact. In presence of insulin, IRs reconstituted into single nanodiscs. All results were consistent with those obtained with IR reconstituted into larger nanodiscs (**SI Figs. 7 - 10**).

Little is known about how IRs transmit signals across the membrane, giving rise to conflicting models (8,11–15). In one model, the IR TMDs interact with each other in the inactive state and insulin binding would pry them apart (15). In a contrasting model, the ECDs keep the TMDs separate in the inactive state and insulin binding would release this inhibition and allow the TMDs to interact (14). It was shown that tryptic removal of the ECDs results in constitutive kinase activity (16), thus implying that the ECDs prevent autophosphorylation of the TKDs in the non-activated receptor, most likely by restraining the TMDs, and hence the associated TKDs, from interacting with each other. A similar model was previously suggested for the human EGF receptor (17) and our results unambiguously support this model also for the IR (**Fig. 2e**). Ligand binding induces a large conformational change in the IR ECDs, releasing the constraint on the TMDs and allowing them to come together. As a result, the cytoplasmic TKDs can interact with each other, autophosphorylate and initiate the signalling cascade. Indeed, some averages of the receptor in the T-shaped conformation show a strong globular density (arrow in Fig. 2c) that is not seen in averages of the receptor in the U-shaped conformation, suggesting that the TKDs in these averages may be dimerized and more restricted in their localization.

Given the rearrangements of the IR TMDs, the lipid environment likely plays an important role in receptor activation and regulation, and a better understanding of insulin signal propagation across the membrane will greatly aid the structure-guided development of drugs.

## EXPERIMENTAL PROCEDURES

### Expression and purification of full-length IR

cDNA for human insulin receptor isoform A was cloned into a pTT6 vector as described(18), resulting in a construct consisting of the human IRA followed by a C-terminal human rhinovirus 3C protease cleavage site and a tandem purification (TAP) tag. The receptor was transiently expressed in suspension-adapted FreeStyle™ 293-F Cells maintained in protein-free FreeStyle™ 293 Expression Medium (Invitrogen) as described (18). The recombinant protein was purified in batch mode in a single-affinity chromatography step using only the immunoglobulin G (IgG)-binding domain of the *Staphylococcus aureus* protein A (ProtA)(18,19) portion of the TAP tag. The following modifications were introduced: the pH of all buffers was adjusted to 7.5 and, in an additional last wash step, the IgG Sepharose 6 Fast Flow resin (GE Healthcare) was washed with 2 column volumes (CV) of 1% (w/v) 3-((3-cholamidopropyl) dimethylammonio)-1-propanesulfonate (CHAPS, Glycon Biochemicals GmbH, Luckenwalde, Germany) and 20% (w/v) glycerol in 50 mM HEPES, pH 7.5, 150 mM NaCl. The resin was then equilibrated with 20 CV elution buffer (1% CHAPS and 8% glycerol in 50 mM HEPES, pH 7.5, 150 mM NaCl), and the protein was cleaved overnight at 4°C by incubation with glutathione S-transferase (GST)-tagged human rhinovirus 3C protease (50 μL per mL IgG beads). The IR was eluted with elution buffer in 750-μL fractions. Co-eluting 3C protease was removed with Glutathione Sepharose 4B (GE Healthcare) resin, and the IR concentration was estimated using a molar extinction coefficient of 377,585 M^-1^ cm^-1^ as calculated by ProtParam (ExPASy) assuming 42 of the total 94 cysteins are forming disulfide bonds (7). For quality controls, analytical size exclusion chromatography was carried out using a small Superose 6 PC 3.2/30 column with IR elution buffer at a flow rate 0.05ml/min at room temperature, followed by SDS page of the collected fractions and silver staining. Purified IR was immediately reconstituted into nanodiscs.

### Purification of MSP1E3D1 and MSP1D1

Plasmids encoding membrane scaffold proteins (MSP) pMSP1E3D1 (plasmid #20066) and pMSP1D1 (plasmid #20061) were purchased from Addgene. Both MSP variants were produced in *E. coli* BL21Gold(DE) (Agilent/Strategene) and purified as described (20), with the exceptions that pre-packed His-Trap HP columns (GE Healthcare) were used for immobilized metal ion affinity chromatography and that an additional wash step with 32 mM cholate was introduced to remove residual bacterial lipids from MSP. The His tag was not removed.

### Reconstitution of IRs into nanodiscs

1,2-dioleoyl-sn-glycero-3-phosphocholine (DOPC), N-stearoyl-D-erythro-sphingosyl-phosphorylcholine (SM) and cholesterol were purchased from Avanti Polar Lipids (Alabama, USA). All lipid stocks were quantified on a regular basis to correct for solvent evaporation by phosphate assay for phospholipids (21), Amplex Red Cholesterol Assay Kit (Invitrogen) and controlled for their stability by thin-layer chromatography. DOPC, SM, and cholesterol were mixed at a molar ratio of 8:1.5:0.5, dried under a stream of nitrogen, and left under vacuum overnight to remove residual solvent. Liposomes were formed by addition of reconstitution buffer (20 mM HEPES, pH 7.5, 100 mM NaCl) with vigorous shaking at 54°C, followed by ten freeze-thaw cycles and extrusion through a 100-nm pore-size polycarbonate filter. Liposomes were solubilized by addition of cholate (Alfa) to a final concentration of 18 mM and sonication.

IRs were reconstituted into nanodiscs in the presence or absence of human insulin (Sigma, #I2643). For reconstitutions in the presence of insulin, 0.3 μM IR in 50 mM HEPES, pH 7.5, 150 mM NaCl, 1% CHAPS, 8% glycerol was incubated with 1 μM insulin for 1 h on ice. Prior to usage, insulin was dissolved to a concentration of 180 μM in 5 mM HCl. Insulin was added to the IR reconstitution mix to a final concentration of 1 μM. Even though the pH of the solution did not change, the same amount of 5 mM HCl was added to the unliganded receptor to rule out potential effects of HCl on IR. For nanodisc formation, 0.3 nmol liganded or unliganded IR was incubated with 200 nmol cholate-solubilized lipids and 10 nmol MSP1E3D1 or MSP1D1 for 30 min in the presence of 13.2 mM CHAPS at 25°C with gentle shaking at 300 rpm. The reconstitution mix was centrifuged at 20,000 g for 20 min at 4°C, and the supernatant was extensively dialysed against at least a 1000-times excess of 20 mM HEPES, pH 7.5, 100 mM NaCl for 26 h at 4°C with 3 buffer exchanges followed by a fourth exchange to 25 mM HEPES, pH 7.5, 150 mM NaCl for 18h. Dialysis was carried out in Spectra/Por^®^ 7 Standard RC pre-treated dialysis tubings (Spectrum Labs) with a molecular weight cut-off of 10 kDa (for the first 18 h) and 50 kDa (until the end of the dialysis). Because of the aggregation propensity of non-reconstituted MSP, aggregates were removed by centrifugation at 150,000 g for 45 min at 4°C. The supernatant was snap-frozen in liquid nitrogen and stored at -80°C. IR-containing nanodiscs were separated from empty nanodiscs by size-exclusion chromatography using a Superose 6 10/300 GL column (GE Healthcare) with 25 mM HEPES, pH 7.5, 150 mM NaCl, and only the peak fraction was collected.

For EM studies of IR reconstituted into nanodiscs with MSP1E3D1, the peak fraction was split into four aliquots. One aliquot was imaged directly by negative-stain EM, while the other aliquots were incubated with varying amounts of insulin (final concentrations of 0.8 nM, 12 nM or 1 μM) for 1 h at 4°C prior to imaging. For IR reconstituted into nanodiscs with MSP1D1, the peak fraction was split into halves. One half was imaged directly by negative-stain EM, while the other half was incubated with 1 μM insulin (final concentration) for 1 h at 4°C prior to imaging.

The concentration of IR in nanodiscs was estimated by UV spectroscopy at 280 nm with a molar extinction coefficient of 438,030 M^-1^ cm^-1^ for samples reconstituted with MSP1E3D1 and 420,445 M^-1^ cm^-1^ for samples reconstituted with MSP1D1 as calculated with ProtParam (ExPASy) assuming that 42 of the 94 cysteins form disulfide bonds.

### *In vitro* phosphorylation assays

A saturating insulin concentration (200 nM) was added to purified CHAPS-solubilized IR or nanodisc-embedded IR and allowed to bind for 1 h on ice. Phosphorylation was carried out with 0.25 mM ATP in 25 mM HEPES, pH 7.5, 150 mM NaCl, 10 mM MgCl_2_, 1 mM MnCl_2_ in a final volume of 25 μl and incubated for 12 min at 25°C. For solubilized IR, the CHAPS concentration was kept at the critical micellar concentration of 0.6% (w/v). The reaction was stopped by addition of SDS sample buffer supplemented with 2.5 mM EDTA. The reaction mixtures were subjected to SDS-PAGE, and IR TKD phosphorylation was assessed by Western blotting.

### SDS-PAGE and Western blots

Electrophoresis was carried out with precast 4-12% Bis-Tris gels or 3-8% Tris-Acetate gels or NativePAGE™ 3-12% Bis-Tris gels (Invitrogen) using the corresponding commercial running buffers (Invitrogen). Proteins were stained with Coomassie blue or silver stain or were transferred to PVDF membranes for immunodetection. Membranes were blocked overnight with 5% (w/v) non-fat dry milk powder in TBST (0.1% (v/v) Tween20 in 20 mM Tris, pH 7.4, 150 mM NaCl) supplemented with phosphatase inhibitors (1 μM NaVO_4_, 20 μM β-glycerophosphate). Membranes were probed with antibodies against phospho-IRβ (Tyr-1150/1151; Cell Signaling Technology Cat# 3024, RRID:AB_331253); 1:1,000) or against total IRβ C terminus (Cell Signaling Technology Cat# 3020, RRID:AB_2249166;1:2,000). Secondary antibodies, goat anti-mouse or goat anti-rabbit, conjugated to horseradish peroxidase (BioRad) were used at a 1:10,000 dilution, and detection was performed using SuperSignal™ West Femto Maximum Sensitivity Substrate electrochemiluminescence substrate (Pierce) with a CCD imager (ImageQuant LAS, Amersham). For uncropped and original Native and SDS Gels as well as Western Blots used in the manuscript see SI Fig. 13 and 14.

### Specimen preparation and electron microscopy

An aliquot (3.5 μL) of full-length IR reconstituted into nanodiscs was adsorbed to a glow-discharged 200-mesh copper grid covered with a thin carbon-coated plastic film and negatively stained with 0.75% (w/v) uranyl formate as described (22). Images were collected using an XR16L-ActiveVu CCD camera (AMT, Woburn, MA) on a Philips CM10 electron microscope (FEI, Hillsboro, OR) operated at an acceleration voltage of 100 kV. The calibrated magnification was 41,513× (nominal magnification of 52,000×), yielding a pixel size of 2.65 Å at the specimen level. The defocus was set to –1.5 μm.

### Image processing

Images of nanodisc-embedded IR showed a very heterogeneous particle population. Therefore, for further image processing, particle selection focused on particles that showed one nanodisc with an additional density extending from them or two nanodiscs connected by a density. For each sample, ~10,000 particles (the exact number of images and selected particles are summarized in **SI Table 1**) were manually selected using the e2boxer.py command of the EMAN2 software package (23) and windowed into 180×180-pixel images. After image normalization and particle centring, the particle images were classified into 200 groups using *Κ*-means classification procedures implemented in SPIDER (24). For the iterative stable alignment and clustering (ISAC) algorithm (25) implemented in SPARX(26), the particle images were reduced to 76×76 pixels and subjected to several ISAC generations specifying 50 images per group and a pixel error threshold of 0.7 (the number of ISAC generations and resulting number of averages for each sample are summarized in **SI Table 1**).

To estimate the approximate number of receptors adopting a particular conformation, we used the *K*-means classification into 200 groups and visually assigned the averages to show 1) a U-shaped IR reconstituted into one nanodisc (1U), 2) a U-shaped IR reconstituted into two nanodiscs (2U), 3) a T-shaped IR reconstituted into one nanodisc (1T), 4) a T-shaped IR reconstituted into two nanodiscs (2T), 5) a II-shaped IR (II) (we only observed IR in this conformation associated with one nanodisc), and 6) an L-shaped IR (L) (we only observed L-shaped IR associated with one nanodisc, consistent with the assumption that these particles represent IR monomers). For each set of averages, the particles in the classes assigned to one of these 6 classes were summed up and are listed in **SI Table 2**. Many of the averages could not be unambiguously assigned to one of the six classes. These classes (X) were considered uninterpretable and were excluded from the quantification. The assignments of the classes for each experimental condition are denoted on the SI figures showing the 200 SPIDER class averages. In addition, the particles showing the L-shaped non-physiological monomeric IR were not considered when calculating the percentage of dimeric receptors in each conformation (1U, 2U, 1T, 2T or II) listed in **SI Table 3**.

## ACKNOWLEDGEMENTS

The authors acknowledge Karim Fahmy and Jana Oertel, Helmholtz Zentrum Rossendorf Dresden, for initial support with nanodisc production. Financial support was provided by the Deutsche Forschungsgemeinschaft (DFG) TRR83 TP18 (ÜC), the DFG-funded Dresden International Graduate School for Biomedicine and Bioengineering (GS97) (TG) and the German Federal Ministry of Education and Research grant to the German Center for Diabetes Research (DZD e.V.) (ÜC). This research was also supported by a postdoctoral fellowship from the Canadian Institutes of Health Research (KHK) and a Helmsley Postdoctoral Fellowship at The Rockefeller University (to KHK).

## CONFLICT OF INTERESTS

The authors declare no conflict or competing financial interests.

### AUTHOR CONTRIBUTIONS

Ü.C. and T.W. designed the experiments. T.G. and M.G. performed the biochemistry and K.H.K. the EM experiments. T.G., K.H.K., M.G, T.W. and Ü.C. analysed the data and wrote the manuscript.

